# HiC4D: Forecasting spatiotemporal Hi-C data with residual ConvLSTM

**DOI:** 10.1101/2022.09.10.507434

**Authors:** Tong Liu, Zheng Wang

## Abstract

**Motivation:** The Hi-C experiments have been extensively used for the studies of mammalian genomic structures. In the last few years, spatiotemporal Hi-C has significantly contributed to the study of genome dynamic reorganization. However, computationally forecasting spatiotemporal Hi-C data still has not been seen in the literature.

**Results:** We present HiC4D for addressing the problem of forecasting spatiotemporal Hi-C data. We designed and tested a novel network, which is a combination of residual network and convolutional long short-term memory (ConvLSTM), and named it residual ConvLSTM (ResConvLSTM). We evaluated our new method and compared it with other four methods including three outstanding video-prediction methods from the literature: ConvLSTM, spatiotemporal LSTM (ST-LSTM), and simple video prediction (SimVP), and one self-designed naïve network (NaiveNet) as a baseline. We used four different spatiotemporal Hi-C datasets for the blind test, including two from mouse embryogenesis, one from somatic cell nuclear transfer (SCNT) embryos, and one from human embryogenesis. Our evaluation results indicate that ResConvLSTM almost always outperforms the other four methods on four blind-test datasets in terms of accurately reproducing spatiotemporal Hi-C contact matrices at future time steps. Our benchmarks also indicate that all five methods can successfully recover the boundaries of topologically associating domains (TADs) called on the experimental Hi-C contact matrices.

**Availability:** HiC4D is publicly available at http://dna.cs.miami.edu/HiC4D/.

## 1 Introduction

Since the Hi-C technique was introduced (Lieberman-Aiden, et al., 2009), it played a vital role in identifying topologically associating domains (TAD) (Dixon, et al., 2012), discovering A/B compartments (Lieberman-Aiden, et al., 2009), and detecting DNA loops (Rao, et al., 2014). The genome-wide high-resolution Hi-C data were widely used in various studies, such as reconstructing chromatin three-dimensional (3D) structures (Nagano, et al., 2017), predicting DNA methylation (Wang, et al., 2016), investigating neural development (Bonev, et al., 2017), and exploring Xist transcript mechanism (Engreitz, et al., 2013).

In recent years, with the coming of the four-dimensional (4D) nucleome project (Dekker, et al., 2017), the dynamics of chromatin architectures during a specific nuclear or cellular process attracted much attention. Various spatiotemporal Hi-C experiments were conducted for investigating different nuclear or cellular developments, such as cardiogenesis (Bertero, et al., 2019), neural differentiation (Bonev, et al., 2017), B cell differentiation (Vilarrasa-Blasi, et al., 2021), B cell reprogramming (Stadhouders, et al., 2018), and embryogenesis of various species including mouse (Chen, et al., 2020; Du, et al., 2017; Ke, et al., 2017), human (Chen, et al., 2019), drosophila (Hug, et al., 2017), Xenopus tropicalis (Niu, et al., 2021), and zebrafish (Wike, et al., 2021). These spatiotemporal Hi-C studies revealed transitions, emergences, and reorganizations of chromatin architectures, and the captured spatiotemporal Hi-C data are an excellent source for exploring the relationships between gene expression and dynamics of genome reprogramming or development.

Long short-term memory (LSTM) (Hochreiter and Schmidhuber, 1997) as a special recurrent neural network (RNN) for capturing long-term dependencies has been well studied and widely used in various areas. The Convolutional LSTM (ConvLSTM) (Xingjian, et al., 2015), a combination of convolutional and LSTM operations, is capable of considering spatial correlations compared with fully connected LSTM (FC-LSTM). The novel architecture named spatiotemporal long short-term memory (ST-LSTM) (Wang, et al., 2017; Wang, et al., 2022) included a new spatiotemporal memory state based on ConvLSTM for learning spatial and temporal representations at the same time. Alphafold (Senior, et al., 2019) used one-dimensional (1D) ConvLSTM for predicting protein structures. Hi-C-LSTM (Dsouza, et al., 2022) used LSTM for learning low-dimensional latent representations of a Hi-C contact matrix.

RNNs based on ConvLSTMs were extensively used in video prediction. The neural networks for video prediction can be roughly split into two big categories (Gao, et al., 2022): (1) RNNs by stacking multiple ConvLSTM-based layers and (2) a spatial encoder and a spatial decoder that are usually implemented as convolutional layers together with a temporal learning part between them. The temporal-learning part in the middle may be RNN, a transformer, or convolutional networks (ConvNet).

One of the newest video-prediction methods that is worth mentioning is the simple video prediction (SimVP) model, which was built completely by convolutional layers (Gao, et al., 2022) and achieved state-of-the-art performance over five commonly used datasets for the task of video prediction. However, when it comes to the problem of forecasting spatiotemporal Hi-C data, we can hardly find a tool in the literature. There is a computational method 4DMax (Highsmith and Cheng, 2021), which takes spatiotemporal Hi-C data as input for predicting chromatin organizations. It can interpolate Hi-C contact maps between two given time points from its predicted 4D models, but it cannot forecast spatiotemporal Hi-C at future time steps.

In this paper, we present HiC4D for forecasting spatiotemporal Hi-C data. Since residual ConvNet (ResNet) (He, et al., 2016) was extremely successful in building deeper networks, we newly designed and implemented a residual ConvLSTM and named it ResConvLSTM, which is a novel network making ConvLSTMs have more layers and with better learning abilities. In total, we benchmarked five different methods including ConvLSTM, ResConvLSTM, ST-LSTM, SimVP, and a naïve network. Our blind test indicates that our ResConvLSTM almost always outperforms the other four methods on four different spatiotemporal Hi-C datasets.

## 2 Material and methods

### 2.1 Spatiotemporal Hi-C datasets

We used four different spatiotemporal Hi-C datasets in this study. These datasets were of a varying number of time steps and read depths for each time point. The first one (Du, et al., 2017) captured Hi-C data in preimplantation embryos at the following stages: gametes (sperm and MII oocyte), pronuclear stage 5 (PN5) zygotes, early 2-cell, late 2-cell, 8-cell, inner cell masses (ICM), and mouse embryonic stem cells (mES). We chose the Hi-C data of the last six development stages as our first spatiotemporal Hi-C dataset (Fig. 1A and Table 1). We downloaded all of the valid Hi-C read pairs in the format of “allValidPairs” from Gene Expression Omnibus (GEO) under accession number GSE82185. To balance sequencing depths among different time steps, we first obtained long-range (> 20-kb) intra-chromosomal read pairs for each stage, and then downsampled read pairs to 115 million for each stage (Table S1).

**Fig. 1.**
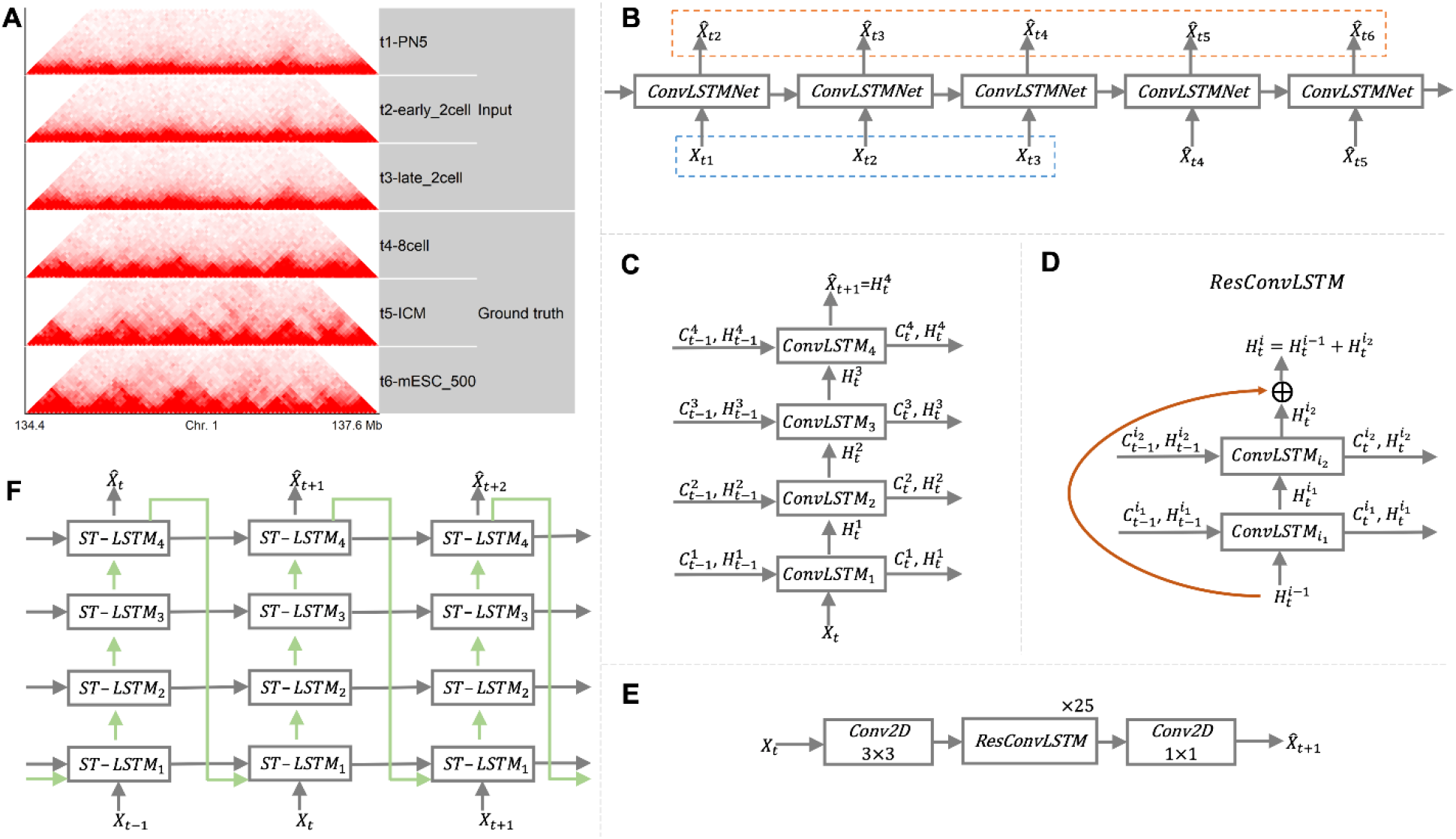
(A) An example of spatiotemporal Hi-C data with six time-steps from dataset 1. (B) The unrolled RNN architecture of a typical next-frame prediction method. The dashed blue box highlights the three time-steps as input. The contents of the dashed orange box consist of the five next-frame reconstructions. (C) A 4-layer ConvLSTM network at time *t*. (D) One block of ResConvLSTM containing two ConvLSTM layers. (E) The architecture of ResConvLSTM network for next-frame prediction at time *t*. (F) A 4-layer ST-LSTM network at times *t* − 1, *t*, and *t* + 1. The blue arrows denote the flows of spatiotemporal memory states.

**Table 1.** Spatiotemporal Hi-C datasets.

| ID | Species | No. of time steps (we used) | System | Genome build | References | Details |
| --- | --- | --- | --- | --- | --- | --- |
| 1 | Mouse | 8 (6) | Embryogenesis | mm9 | (Du, et al., 2017) | Table S1 |
| 2 | Mouse | 8 (6) | Embryogenesis | mm10 | (Ke, et al., 2017) | Table S2 |
| 3 | Mouse | 13 (6) | SCNT embryos | mm10 | (Chen, et al., 2020) | Table S3 |
| 4 | Human | 6 (5) | Embryogenesis | hg19 | (Chen, et al., 2019) | Table S4 |

The second dataset (Ke, et al., 2017) is very similar to the first one, and it captured Hi-C data of mouse gametes (sperm and MII oocyte) and early embryos (Table 1), including 2-cell, 4-cell, 8-cell, embryonic day (E)3.5, and E7.5 stages. As in dataset 1, we only used the last six stages in this study. We downloaded raw Hi-C reads from Genome Sequence Archive (GSA) under accession number PRJCA000241. We mapped the raw reads to reference genome (mm10) using Juicer (Durand, et al., 2016), used the read pair file “merged_nodups.txt” to obtain long-range (> 20-kb) intra-chromosomal read pairs, and finally downsampled read pairs to 62 million for each stage (Table S2).

The third dataset (Chen, et al., 2020) captured spatiotemporal Hi-C of somatic cell nuclear transfer (SCNT) embryos (Table 1). It included 13 different stages of reconstructed embryos. Considering the lower sequencing depths of some stages and being largely consistent with the time steps of dataset 1, we only used six stages (Table S3). We downloaded the Hi-C read pairs under allValidPairs from GEO with accession number GSE146001, obtained long-range intra-chromosomal, and downsampled read pairs (33 million) as above (Table S3).

The last dataset (Chen, et al., 2019) contains six stages of spatiotemporal Hi-C during human embryogenesis including sperm, 2-cell, 8-cell, morula, blastocysts, and six-week-old embryos. We used the last five stages (Table 1) in our experiment. We downloaded the raw reads from GSA under accession number CRA000852. We mapped the raw reads to the reference genome (hg19) using Juicer (Durand, et al., 2016), filtered the read pair file “merged_nodups.txt” to obtain long-range (> 20-kb) intra-chromosomal read pairs, and finally downsampled read pairs to 14.5 million for each stage (**Table S4**).

### 2.2 HiC4D overview

The training data were extracted from dataset 1. For all chromosomes from 1 to X, we extracted validation data on chromosome 19, two chromosomes (i.e., 2 and 6) were left for the blind test, and the rest chromosomes were used for generating training data. The other three datasets were left for a blind test. Considering read depths, we only focused on the resolution of 40 kb in this study. The samples for each chromosome were generated along the diagonal of its 2D raw contact matrix with a sliding window of size 50 × 50 and a step of size 3 bins. The samples were concatenated as an *n* × *t* × 1 × 50 × 50 five-dimensional tensor, where *n* is the total number of samples and *t* is the number of time steps. The main purpose of this study is to use the Hi-C data of the first three time-steps (*t*_1_, *t*_2_, and *t*_3_) as input to predict the corresponding Hi-C data of the last three time-steps (*t*_4_, *t*_5_, and *t*_6_).

We tested two types of frame prediction methods: next-frame and 3-step ahead. The next-frame methods consist of ConvLSTM, our newly designed ResConvLSTM, and ST-LSTM. The 3-step ahead methods include SimVP and NaiveNet. These five methods were trained with the same data. The best models for the blind test were the ones that achieved the best performance on validation data. Since one pixel may be predicted more than one time, its final prediction is the average value of all predictions.

After obtaining predictions for each testing chromosome at the future time steps, we evaluated each method mainly by quantifying the similarity scores between ground truth and predictions for each time step.

### 2.3 Next-frame method

The input of next-frame methods *X*_*t*_ at time *t* is an *n* × 1 × 50 × 50 four-dimensional tensor. After passing through the network, we obtain the output 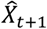, which is thought of as the reconstruction of *X*_*t*+1_ (Fig. 1B). Since our input time steps are *t*_1_, *t*_2_, and *t*_3_, the fourth and the following *X*_*t*_ are directly from the output of their previous time step (Fig. 1B). The loss is calculated between *X*_*t*_ and reconstructed 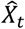, where *t* is from two to six. Technical details of three next-frame methods (ConvLSTM, ResConvLSTM, and ST-LSTM) will be presented in the following three subsections.

#### 2.3.1 Convolutional LSTMs

The first convolutional LSTM (ConvLSTM-1) we implemented and benchmarked in this study is the same as the one published in the paper that first applied convolutional operation to LSTM (Xingjian, et al., 2015). The following equations describe a typical *l*-th ConvLSTM-1 layer at time *t* with the following three inputs: the input data *X*_*t*_, the last memory cell state 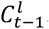, and the last hidden state 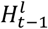.

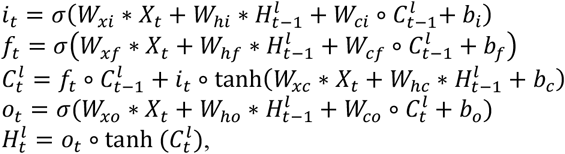

where ∗ denotes the convolution operator, *σ* denotes the sigmoid function, ∘ denotes the Hadamard product, and *W* and *b* are the weight and bias parameters that need to be learned. Since there are various LSTM variants, we added two more ConvLSTMs (i.e., ConvLSTM-2 and ConvLSTM-3). When calculating *i*_*t*_, *f*_*t*_, and *o*_*t*_, we removed the Hadamard products in ConvLSTM-2 and replaced all the Hadamard products with convolutional operations in ConvLSTM-3 (see supplementary materials for details). The other parts for the three ConvLSTMs remain the same. We only used one of them for the blind test, and the evaluation results for the three ConvLSTM variants are shown in the Results section. We built the ConvLSTM networks by simply stacking multiple ConvLSTM layers (Fig. 1C).

#### 2.3.2 Residual convolutional LSTM

We designed residual ConvLSTM (ResConvLSTM), which is a combination of residual ConvNet and ConvLSTM (Fig. 1D). The *i*-th ResConvLSTM block shown in Fig. 1D contains two ConvLSTM-1 layers (i.e., 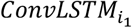 and 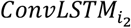). The final output 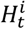 of this block at time *t* is equal to

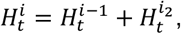

where the input 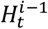 is the output of the previous ResConvLSTM block and the updated hidden state 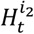 is the output of 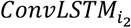. The skip connection can make the output 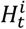 directly link to the input 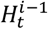. The final ResConvLSTM network we used for the blind test is shown in Fig. 1E. It contains two 3 × 3 2D convolutional layers for increasing and reducing hidden channels, and the middle part is built by stacking 25 ResConvLSTM blocks.

#### 2.3.3 Spatiotemporal LSTM

The spatiotemporal LSTM (ST-LSTM) introduced a novel spatiotemporal memory state (Wang, et al., 2017). It can flow in both bottom-up and top-down directions, whereas the memory cell state and the hidden state can only flow in horizontal and bottom-up directions, respectively. The following equations describe a ST-LSTM layer at time *t*:

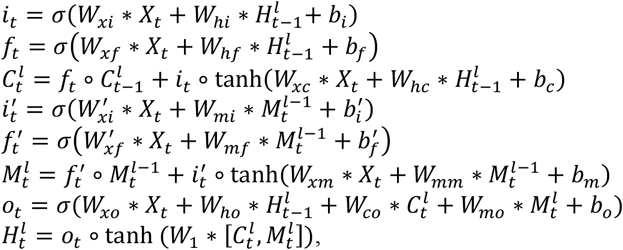

where 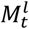 is the spatiotemporal memory state for the *l*-th layer at time *t* and *W*_1_ is 1 × 1 2D convolutional operation for reducing the hidden dimension. The process of generating 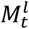 is very similar to deriving 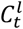. We implemented the final ST-LSTM networks by stacking multiple ST-LSTM layers (Fig. 1F).

### 2.4 The 3-step ahead methods: SimVP and NaiveNet

As its name suggests, the 3-step ahead methods mean that we predict three future frames in one prediction. We used the same network of SimVP as described in (Gao, et al., 2022). The batch size and the hidden dimension were set to 32 and 128, respectively. The other hyperparameters were with defaults. We designed a naïve 3D convolutional neural network (NaiveNet) as a baseline. The NaiveNet (Fig. S1) contains three 3D convolutional layers. Each of the first two layers is followed by Group normalization (number of groups set to 2) (Wu and He, 2018) and LeakyReLU (negative slope set to 0.2).

The model of NaiveNet for the blind test was trained with the following hyperparameters: batch size 32, hidden dimension 128, and kernel sizes 7 for the two spatial dimensions (height and width) and 1 for the temporal dimension. The input of 3-step ahead methods is an *n* × 3 × 1 × 50 × 50 five-dimensional tensor, where 3 denotes the first three time-steps (i.e., *t*_1_, *t*_2_, and *t*_3_). The output is still an *n* × 3 × 1 × 50 × 50 five-dimensional tensor, but here 3 denotes the last three time-steps (i.e., *t*_4_, *t*_5_, and *t*_6_). The loss is calculated between the output and the ground truth at the last three time-steps.

### 2.5 Implementation details

All models were implemented in PyTorch (Paszke, et al., 2019). Unless otherwise stated, the loss function for all the models is mean squared error (MSE), and the optimizer for training all models is Adam (Kingma and Ba, 2014) with a learning rate set to 0.0001. All models were trained on an NVIDIA A100 GPU with 40 GB memory.

### 2.6 Evaluation metrics

The main evaluation strategy is by quantifying similarity or reproducibility between ground-truth and predicted Hi-C contact matrices at the future time steps. The two metrics we used are Pearson correlation coefficients at each genomic distance and stratum-adjusted correlation coefficient (SCC) from HiCRep (Yang, et al., 2017). When running HiCRep, we set the smoothing parameter to 5 and the lower and upper bounds of the genomic distance to 400,000 and 1,600,000, respectively. We also calculated insulation scores (Crane, et al., 2015) (see supplementary materials for details) for evaluating the ability to recover TAD boundaries.

## 3 Results

### 3.1 Hyperparameter tuning and model selection

For training ConvLSTM-based models, we tested various hyperparameter combinations (see Table S5). For the three ConvLSTM networks (ConvLSTM-1, ConvLSTM-2, and ConvLSTM-3), the first two achieved a smaller MSE loss (0.0067 and 0.00669) than the last one (0.00684). To select one for the blind test, we further evaluated the three networks using Pearson correlations on chromosome 19 at each genomic distance (Fig. S2). In general, ConvLSTM-1 performs better than the other two. Therefore, the 4-layer ConvLSTM-1 was selected as the representative of ConvLSTM networks for the blind test.

For tuning ResConvLSTM networks, we found three hyperparameter combinations (see Table S5) achieved almost the same best losses (0.00666, 0.00664, and 0.00666). Therefore, we further evaluated the three models using Pearson correlations (Fig. S3) and observed that deeper ResConvLSTM networks perform better, especially at the time *t*_6_. The model with 25 ResConvLSTM blocks (52 layers) was used for the blind test.

For tuning ST-LSTM networks, we trained two more models with the loss function equal to *MSE* + 0.1 × *decouple* (Wang, et al., 2022). The further evaluation results for the model with the smallest MSE loss and the two more models are shown in Fig. S4, indicating that adding decouple loss during training does not provide better performance. Therefore, the 4-layer ST-LSTM model with the smallest MSE loss was selected for the blind test.

We also reported the MSE losses of two 3-step ahead methods (SimVP and NaiveNet) in Table S5. It is obvious that their losses are larger than those of next-frame methods because the latter also consider the reconstruction errors of times *t*_2_ and *t*_3_, which are usually smaller because spatiotemporal Hi-C at times *t*_2_ and *t*_3_ are easier to learn. Taken together, all the benchmarking models for the blind test were trained with the same batch size of 32 and the same hidden dimension of 128 (32 for ResConvLSTM because of GPU memory limitation). In addition, we also borrowed the wide (Szegedy, et al., 2015) and dense (Huang, et al., 2017) concepts from convolutional neural networks onto ConvLSTMs but did not obtain a smaller validation loss (data and details not shown).

### 3.2 Benchmarks on dataset 1

The reproducibility results for the blind test on dataset 1 are shown in Fig. 2. Our newly-designed ResConvLSTM often had higher Pearson correlations in comparison to the other four methods. The performance of SimVP became worse at the long-term time point (*t*_6_). Moreover, ResConvLSTM mostly outperformed the other four methods when the SCC scores for reproducibility were assessed. One exception is that SimVP achieved a bit higher SCC of 0.851 on chromosome 2 at the time *t*_4_ compared to 0.849 of ResConvLSTM. As expected, NaiveNet consistently had the lowest correlations and SCCs. The TAD-recovering results are shown in Fig. 3 and Fig. S5. We first found that almost all predicted Hi-C contact matrices had very similar insulation scores with ground truth (Fig. S5), and even NaiveNet achieved >0.86 correlations at times *t*_4_ and *t*_5_. We then called strong TAD boundaries on ground-truth Hi-C contact matrices using cooltools (https://github.com/open2c/cooltools), plotted average insulation scores around strong TAD boundaries (Fig. 3A), and observed valleys/minima of average insulation scores calculated on the predicted Hi-C contact matrices from each of the five methods, especially for ResConvLSTM, indicating that these methods can successfully recover the strong TAD boundaries. Moreover, we showed some specific TAD-recovering examples at a genomic region in Fig. 3B and Fig. S6, revealing that our newly-designed ResConvLSTM and the other four methods can successfully recover TAD patterns that have not yet been fully established at the input time steps.

**Fig. 2.**
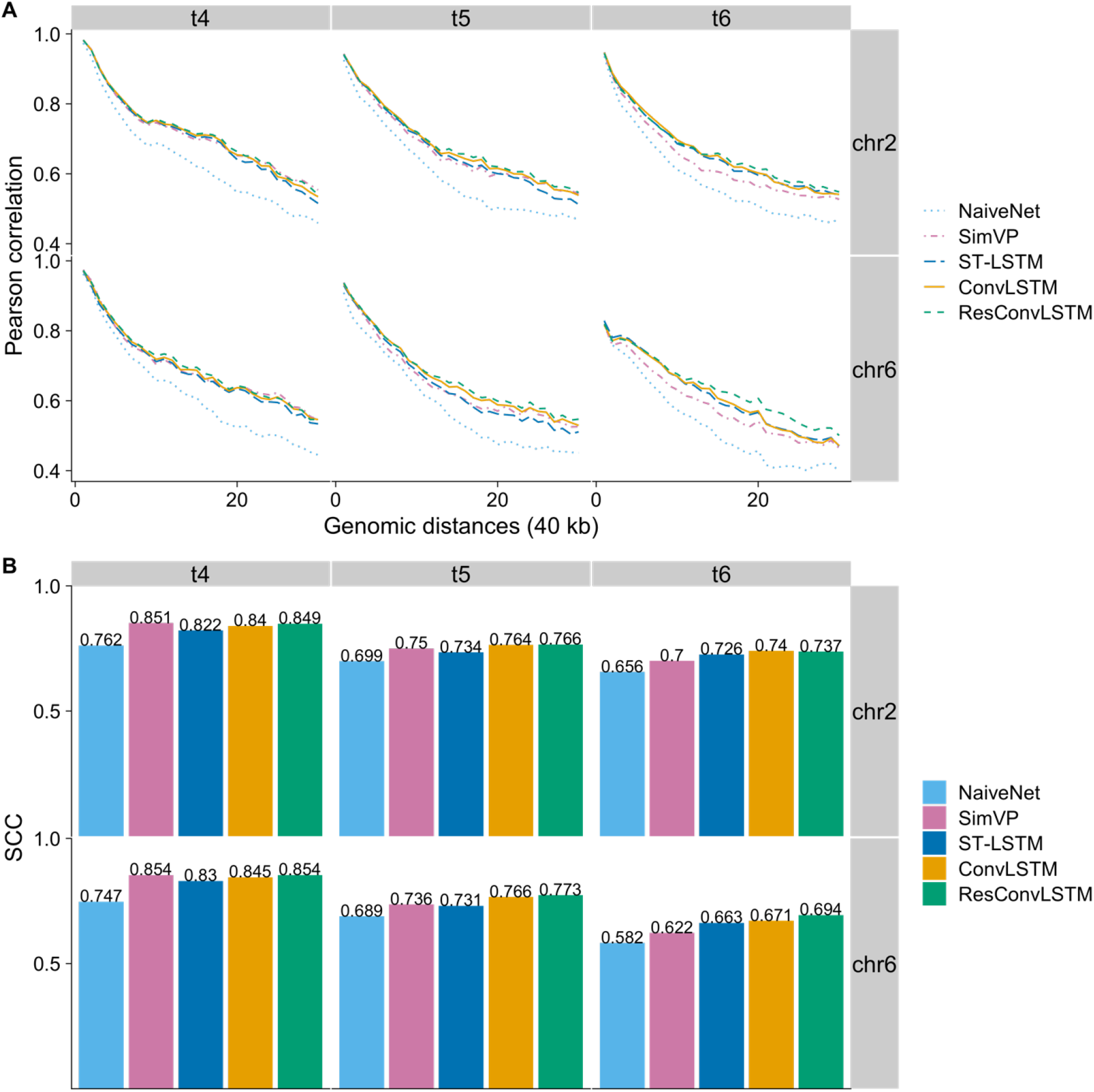
Reproducibility results on dataset 1. (A) Pearson correlations between ground truth and predicted Hi-C contact matrices at each genomic distance. (B) SCC scores between ground truth and predicted Hi-C contact matrices from the five methods.

**Fig. 3.**
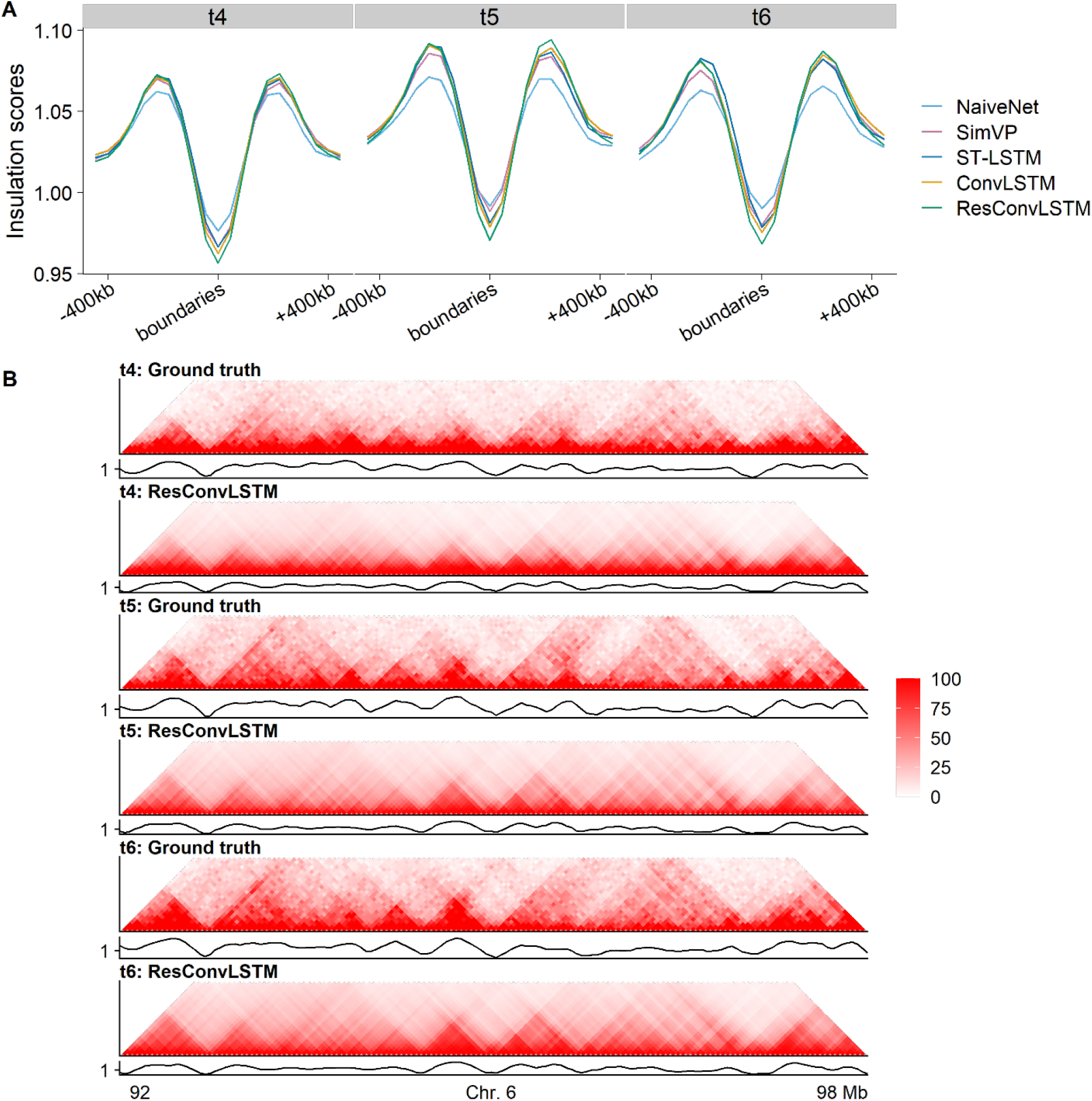
TAD-recovering results on dataset 1. (A) Valleys/minima of insulation scores from predicted Hi-C contact matrices around strong TAD boundaries that were called based on ground-truth Hi-C. (B) Hi-C heat maps and their insulation-score curves for ground truth and ResConvLSTM predictions at the three future time steps (*t*_4_, *t*_5_, and *t*_6_).

### 3.3 Benchmarks on dataset 2

It is worth noting that compared with the other two datasets (3 and 4) dataset 2 is more similar to dataset 1 because of the number of read pairs after downsampling, time-point ranking order, and the fact that they are both from mouse embryogenesis. We report the reproducibility benchmarks in Fig. 4 and TAD-recovering results in Fig. 5 and Fig. S7. At the first two future time-steps (*t*_4_ and *t*_5_), ResConvLSTM often had the highest correlations and SCCs with one exception in dataset 1 that SimVP achieved a bit higher SCC of 0.785 compared to 0.77 for ResConvLSTM on chromosome 2 at time *t*_4_. However, ST-LSTM performed best at time *t*_6_ and was closely followed by our method ResConvLSTM. SimVP somehow performed worse even than NaiveNet at time *t*_6_. As we concluded for dataset 1, the TAD-recovering results (Fig. 5 and Fig. S7) for dataset 2 also indicated that the five methods can successfully recover TAD profiles even though our models were not trained on this dataset.

**Fig. 4.**
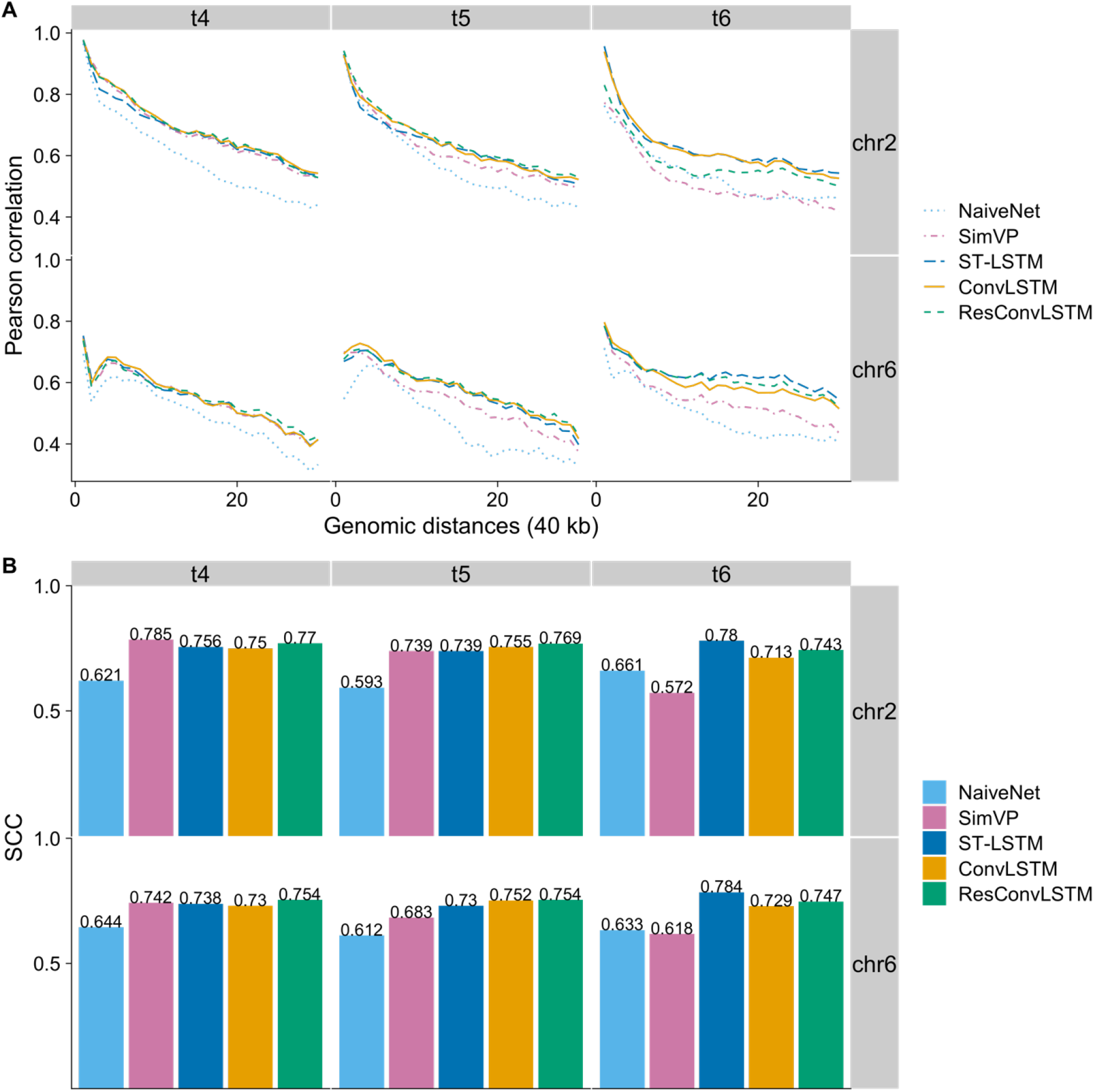
Reproducibility results for dataset 2. (A) Pearson correlations and (B) SCC scores between ground truth and predicted Hi-C contact matrices.

**Fig. 5.**
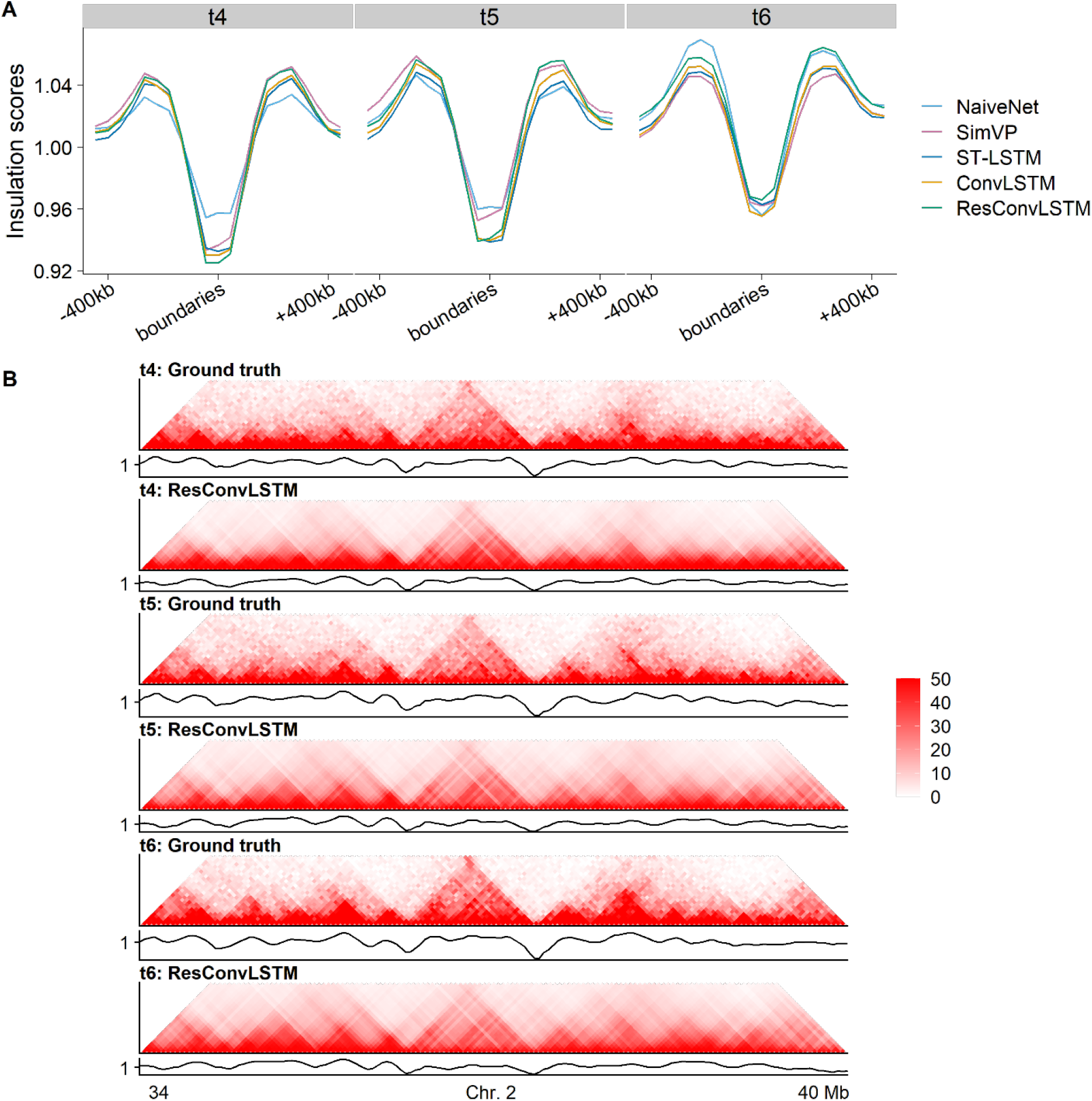
TAD-recovering results for dataset 2. (A) Valleys/minima of insulation scores from predicted Hi-C contact matrices around strong TAD boundaries called on experimental Hi-C. (B) Hi-C heat maps and their corresponding insulation-score curves for ground truth and ResConvLSTM predictions.

### 3.4 Benchmarks on datasets 3 and 4

The third dataset contains 13 time-points, and we only used six of them for matching the time steps of dataset 1. The main difference between dataset 3 and the first two datasets is that spatiotemporal Hi-C data were captured during SCNT embryo development in dataset 3, whereas it was mouse embryos for the first two datasets. The reproducibility benchmarks for dataset 3 are shown in Fig. 6. In general, ResConvLSTM outperforms the other four methods, and the two 3-step ahead methods perform worse.

**Fig. 6.**
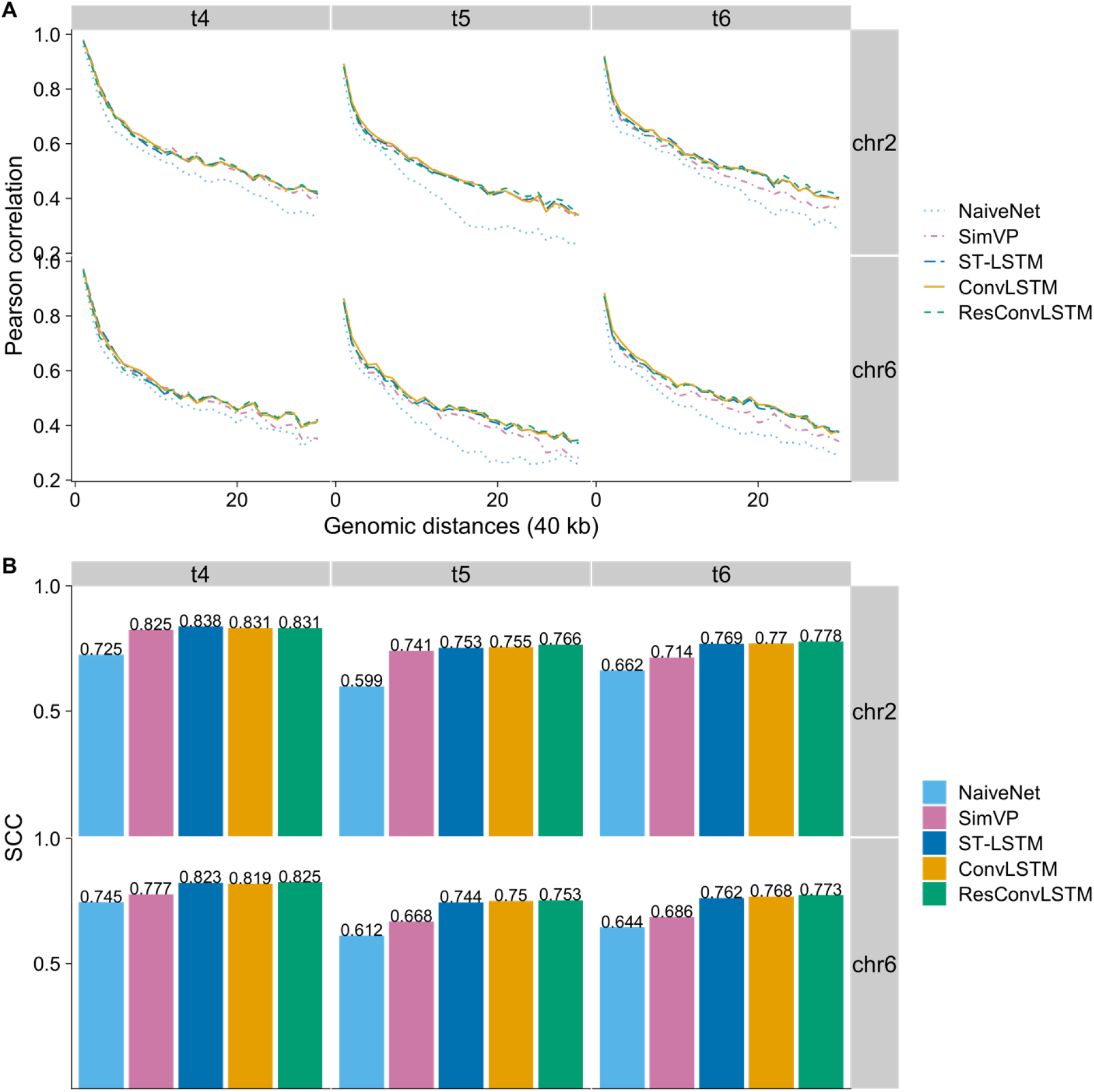
Reproducibility results for dataset 3. (A) Pearson correlations at each genomic distance and (B) SCC scores between ground truth and predicted Hi-C contact matrices.

The spatiotemporal Hi-C data in the last dataset were captured during human embryo development, which help us assess our methods in a different species. The reproducibility results for two future time-steps (*t*_4_ and *t*_5_) are shown in Fig. 7. At time *t*_4_, our ResConvLSTM achieved state-of-the-art performance and is followed by ConvLSTM, ST-LSTM, SimVP, and NaiveNet. At time *t*_5_, ResConvLSTM still performs best, whereas ConvLSTM and SimVP perform worse even than NaiveNet. As we observed from the evaluation results for the first three datasets, it seems that SimVP cannot predict long-term Hi-C matrices as well as predicting short-term ones, whereas our method ResConvLSTM does not have this weakness.

**Fig. 7.**
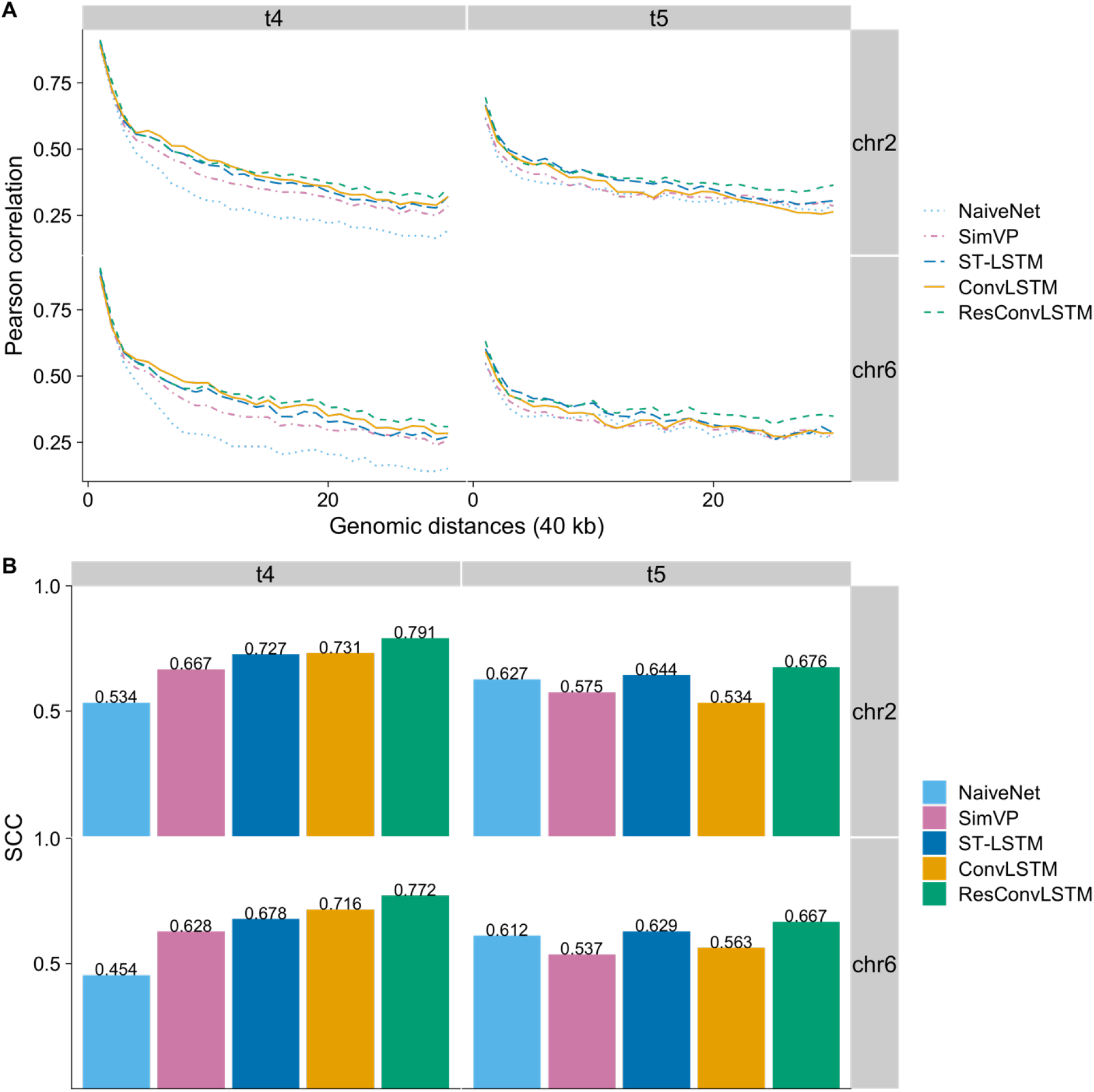
Reproducibility benchmarks for dataset 4. (A) Pearson correlations and (B) SCC scores between ground truth and predicted Hi-C contact matrices at times *t*_4_ and *t*_5_.

## 4 Conclusions

In this paper, we present HiC4D for exploring the forecasting problem of spatiotemporal Hi-C data. We predicted the future three Hi-C frames from the previous three frames with five different video-prediction methods. Specifically, we reimplemented two RNN methods (ConvLSTM and ST-LSTM), introduced a novel method (ResConvLSTM), and used a state-of-the-art method (SimVP) and a NaiveNet as a baseline. These methods were trained with the same data and blindly tested on four different spatiotemporal Hi-C datasets. Our benchmarks indicate that our newly designed method ResConvLSTM almost always outperforms the other four methods across the four datasets by achieving higher reproducibility scores. Our evaluation results also indicate that all five methods can successfully recover TAD boundaries. Together, HiC4D is an effective tool for accurately predicting spatiotemporal Hi-C data.

