## Supplementary materials for "HiC4D: Forecasting spatiotemporal Hi-C data with residual ConvLSTM"

### ConvLSTM-2

$$
\begin{aligned}
i_t &= \sigma(W_{xi} * X_t + W_{hi} * H_{t-1}^{l} + b_i) \\
f_t &= \sigma\left(W_{xf} * X_t + W_{hf} * H_{t-1}^{l} + b_f\right) \\
C_t^{l} &= f_t \circ C_{t-1}^{l} + i_t \circ \tanh(W_{xc} * X_t + W_{hc} * H_{t-1}^{l} + b_c) \\
o_t &= \sigma(W_{xo} * X_t + W_{ho} * H_{t-1}^{l} + b_o) \\
H_t^{l} &= o_t \circ \tanh\,(C_t^{l})
\end{aligned}
$$

### ConvLSTM-3

$$
\begin{aligned}
i_t &= \sigma(W_{xi} * X_t + W_{hi} * H_{t-1}^{l} + W_{ci} * C_{t-1}^{l} + b_i) \\
f_t &= \sigma\left(W_{xf} * X_t + W_{hf} * H_{t-1}^{l} + W_{cf} * C_{t-1}^{l} + b_f\right) \\
C_t^{l} &= f_t \circ C_{t-1}^{l} + i_t \circ \tanh(W_{xc} * X_t + W_{hc} * H_{t-1}^{l} + b_c) \\
o_t &= \sigma(W_{xo} * X_t + W_{ho} * H_{t-1}^{l} + W_{co} * C_t^{l} + b_o) \\
H_t^{l} &= o_t \circ \tanh\,(C_t^{l})
\end{aligned}
$$

### Insulation score

The insulation score (IS) for the $i$-th 40-kb bin is calculated as following:

$$IS_i = \frac{1}{n^2} \sum_{j \in J, k \in K} C(j,k)$$

$$IS_i' = \log_2\left(\frac{IS_i}{IS_{avg}} + 1\right),$$

where $J$ and $K$ are the bin sets of $[i-5, i-1]$ and $[i+1, i+5]$, respectively, $C(j,k)$ is the Hi-C contacts at bin indexes $j$ and $k$, $n$ is equal to 5, and $IS_{avg}$ is the average IS of all bins on the current chromosome. The final IS ($IS_i'$) is the log transformation of $\log_2(x+1)$, where $x$ is the ratio of each bin's IS ($IS_i$) and the average IS ($IS_{avg}$).

**Table S1.** Details of dataset 1 about numbers of valid read pairs at each time step.

| Time ID | Time points | Number of allValidPairs | After filtering (>20kb intra-chr.) | After downsampling |
|---|---|---|---|---|
| t1 | PN5 zygote | 250,033,607 | 136,514,531 | 115,000,000 |
| t2 | Early 2-cell | 238,136,944 | 134,350,801 | |
| t3 | Late 2-cell | 309,906,331 | 170,077,578 | |
| t4 | 8-cell | 290,254,185 | 192,683,578 | |
| t5 | ICM | 221,486,743 | 149,431,330 | |
| t6 | mESC | 158,789,957 | 115,438,254 | |

**Table S2.** Details of dataset 2 about numbers of contact pairs at each time step.

| Time ID | Time points | Number of all contact pairs | After filtering (>20kb intra-chr.) | After downsampling |
|---|---|---|---|---|
| t1 | zygote | 254,205,558 | 62,176,913 | 62,000,000 |
| t2 | 2-cell | 234,656,744 | 64,917,644 | |
| t3 | 4-cell | 656,846,297 | 209,313,083 | |
| t4 | 8-cell | 572,517,384 | 153,136,307 | |
| t5 | E3.5 | 596,167,520 | 178,305,007 | |
| t6 | E7.5 | 1,412,804,902 | 575,229,812 | |

**Table S3.** Details of dataset 3 about numbers of valid read pairs at each time step.

| Time ID | Time points | Number of allValidPairs | After filtering (>20kb intra-chr.) | After downsampling |
|---|---|---|---|---|
| | CC | 137,231,961 | 96,131,399 | 33,000,000 |
| | 0.5hpi | 13,829,278 | 9,385,399 | |
| | 1hpi | 11,321,468 | 8,590,129 | |
| | 1hpa | 8,926,567 | 7,148,171 | |
| | 6hpa | 113,494,879 | 56,164,146 | |
| t1 | 12hpa | 159,766,544 | 78,174,527 | |
| t2 | Early-2-cell | 174,948,961 | 94,369,270 | |
| t3 | Late-2-cell | 89,978,789 | 53,398,992 | |
| | 4-cell | 100,220,085 | 54,414,433 | |
| t4 | 8-cell | 51,205,945 | 33,723,297 | |
| | Morula | 87,366,689 | 57,708,483 | |
| t5 | ICM | 123,165,649 | 80,643,557 | |
| t6 | TE | 165,825,942 | 113,567,317 | |

**Table S4.** Details of dataset 4 about numbers of contact pairs at each time step.

| Time ID | Time points | Number of all contact pairs | After filtering (>20kb intra-chr.) | After downsampling |
|---|---|---|---|---|
| t1 | 2-cell | 40,311,069 | 14,733,663 | 14,500,000 |
| t2 | 8-cell | 69,829,968 | 25,458,239 | |
| t3 | morula | 43,918,745 | 14,557,114 | |
| t4 | blastocyst | 755,470,274 | 310,951,826 | |
| t5 | 6-week | 596,229,576 | 235,384,135 | |

**Table S5.** Validation results of hyperparameter tuning for three next-frame methods (ConvLSTM, ResConvLSTM, and ST-LSTM) and two 3-step ahead methods (SimVP and NaiveNet). The further evaluation results for the highlighted models are shown in Results section.

| Method | Loss | Batch size | Hidden dimension | Kernel size | Number of layers | Validation loss |
|---|---|---|---|---|---|---|
| ConvLSTM-1 | MSE | 32 | 128 | 5 | 4 | 0.00676 |
| | | 16 | 128 | 5 | 8 | 0.00694 |
| | | 16 | 128 | 5 | 12 | 0.00686 |
| | | **32** | **128** | **7** | **4** | **0.00670** |
| | | 32 | 128 | 11 | 4 | 0.00677 |
| | | 32 | 128 | 7 | 2 | 0.00673 |
| | | 32 | 32 | 7 | 4 | 0.00679 |
| ConvLSTM-2 | MSE | 32 | 128 | 5 | 4 | 0.00672 |
| | | **32** | **128** | **7** | **4** | **0.00669** |
| | | 32 | 128 | 7 | 2 | 0.00678 |
| | | 32 | 128 | 9 | 4 | 0.00671 |
| ConvLSTM-3 | MSE | 32 | 128 | 5 | 4 | 0.00684 |
| | | **32** | **128** | **7** | **4** | **0.00687** |
| ResConvLSTM | MSE | 32 | 128 | 7 | 6 | 0.00669 |
| | | **32** | **128** | **7** | **10** | **0.00666** |
| | | 32 | 128 | 7 | 14 | 0.00676 |
| | | **32** | **64** | **7** | **30** | **0.00664** |
| | | 32 | 64 | 5 | 30 | 0.00677 |
| | | **32** | **32** | **7** | **52** | **0.00666** |
| | | 32 | 16 | 7 | 102 | 0.00673 |
| ST-LSTM | MSE | **32** | 128 | **5** | **4** | **0.00667** |
| | | 16 | 128 | 5 | 8 | 0.00683 |
| | | 16 | 128 | 5 | 12 | 0.00682 |
| | | 32 | 128 | 7 | 4 | 0.00676 |
| | | 32 | 128 | 11 | 4 | 0.00675 |
| | MSE + decouple | **32** | **128** | **5** | **4** | **0.00690** |
| | | **32** | **128** | **7** | **4** | **0.00689** |
| SimVP | MSE | 32 | 128 | - | - | 0.00944 |
| NaiveNet | MSE | 32 | 128 | 7 | 3 | 0.01105 |

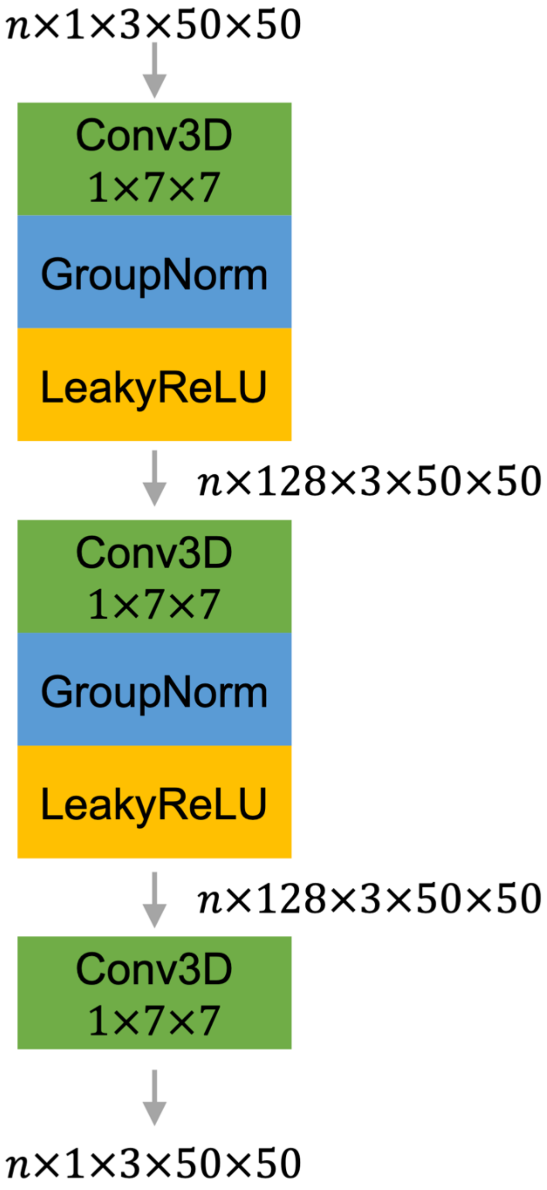


**Fig. S1.** The architecture of the 3-layer NaiveNet. The number of groups for each of the two group normalizations (GroupNorm) is set to 2. The negative slope for each of the two LeakyReLUs is set to 0.2. The padding tuple for each of the three Conv3Ds is (1, 3, 3) for keeping the shape of $3 \times 50 \times 50$. The three input temporal channels are $t_1$, $t_2$, and $t_3$, while the three output temporal channels are corresponding to $t_4$, $t_5$, and $t_6$.

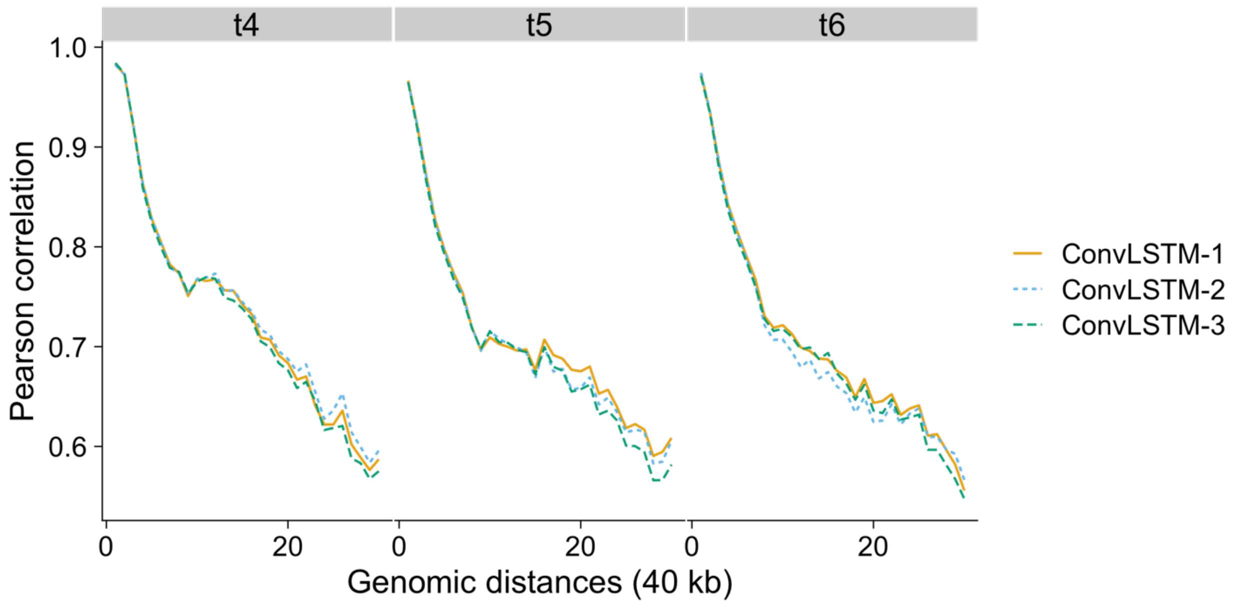


**Fig. S2.** Performances of three ConvLSTM networks on validation data from chromosome 19 at times $t_4$, $t_5$, and $t_6$. Pearson correlations between experimental Hi-C and predicted Hi-C from each of the three ConvLSTMs at each genomic distance. The three ConvLSTMs were trained with the same hyperparameters: the batch size of 32, the kernel size of 5, and the number of layers equal to 4.

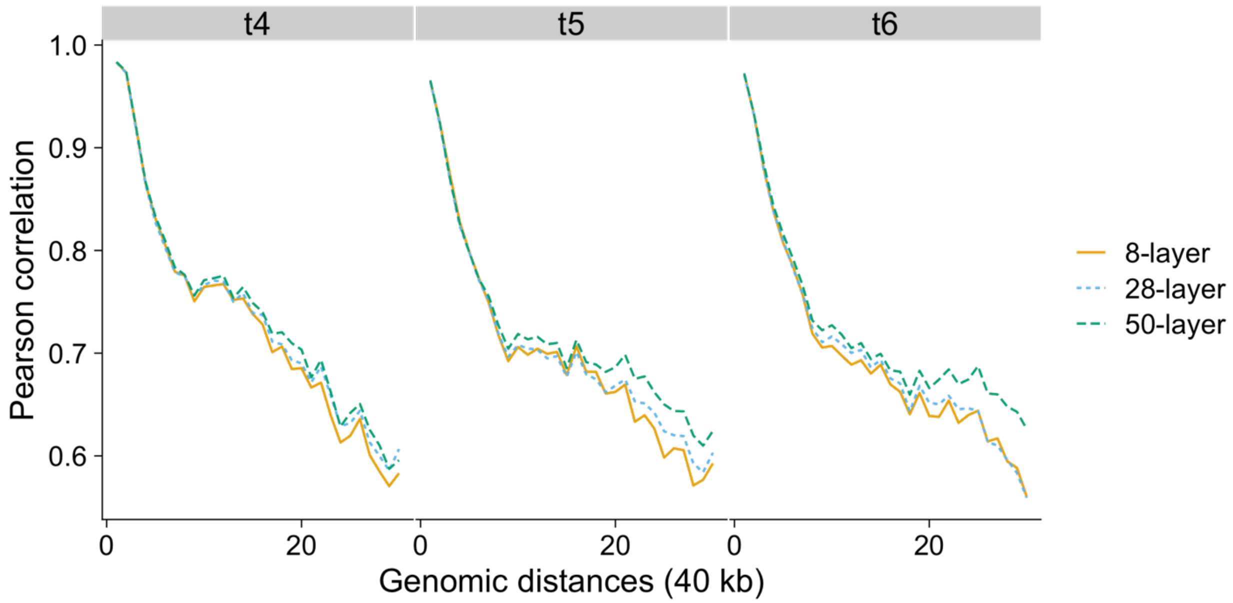


**Fig. S3.** Deeper ResConvLSTM performs better on validation data from chromosome 19. Pearson correlations between experimental Hi-C and predicted Hi-C from each of the three ResConvLSTMs at each genomic distance. The three ResConvLSTM networks were trained with the same kernel size of 7 and the same batch size of 32, but with different hidden dimensions and number of layers because of the limitation of GPU memory. The hidden dimensions of the 8-layer, 28-layer, and 50-layer networks (together with two more convolutional layers) are 128, 64, and 32, respectively.

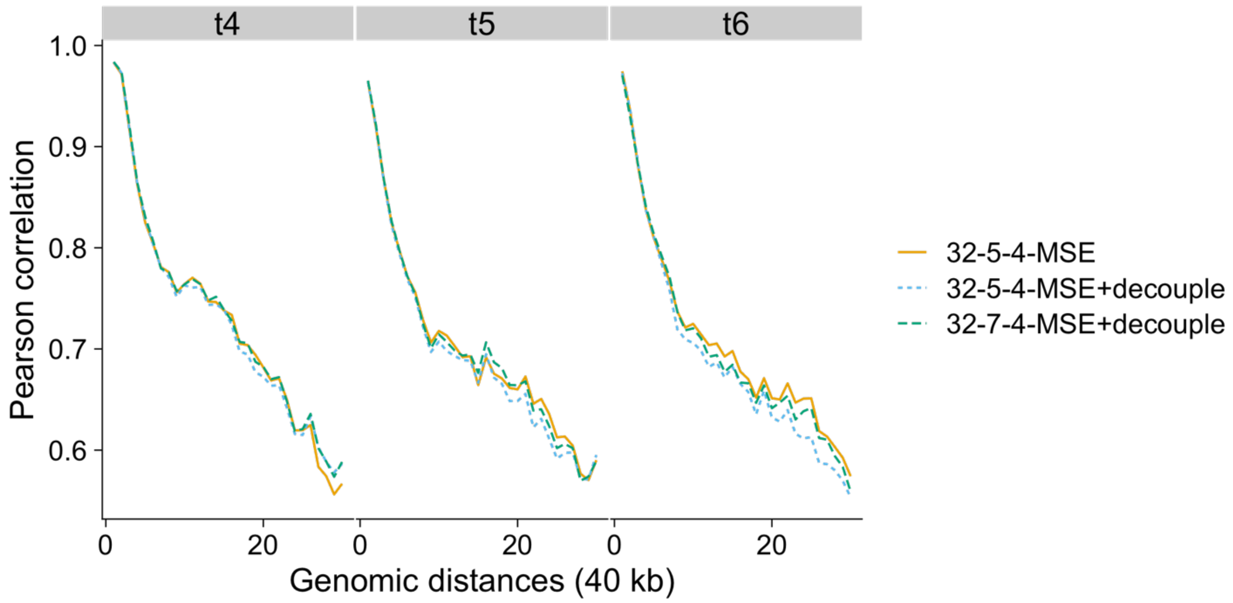


**Fig. S4.** ST-LSTM without decouple loss performs better on validation data from chromosome 19 at time $t_6$. Pearson correlations between experimental Hi-C and predicted Hi-C from each of the three ST-LSTMs at each genomic distance. The three ST-LSTMs were trained with different configurations. The first one was equipped with the batch size of 32, the kernel size of 5, and the number of layers equal to 4 together with only MSE loss. The second one was equipped with the same hyperparameters as the first one but used MSE plus decouple loss. The last one was equipped with the same batch size (32) and the number of layers (4), but with the kernel size of 7 together with MSE plus 0.1 times decouple loss.

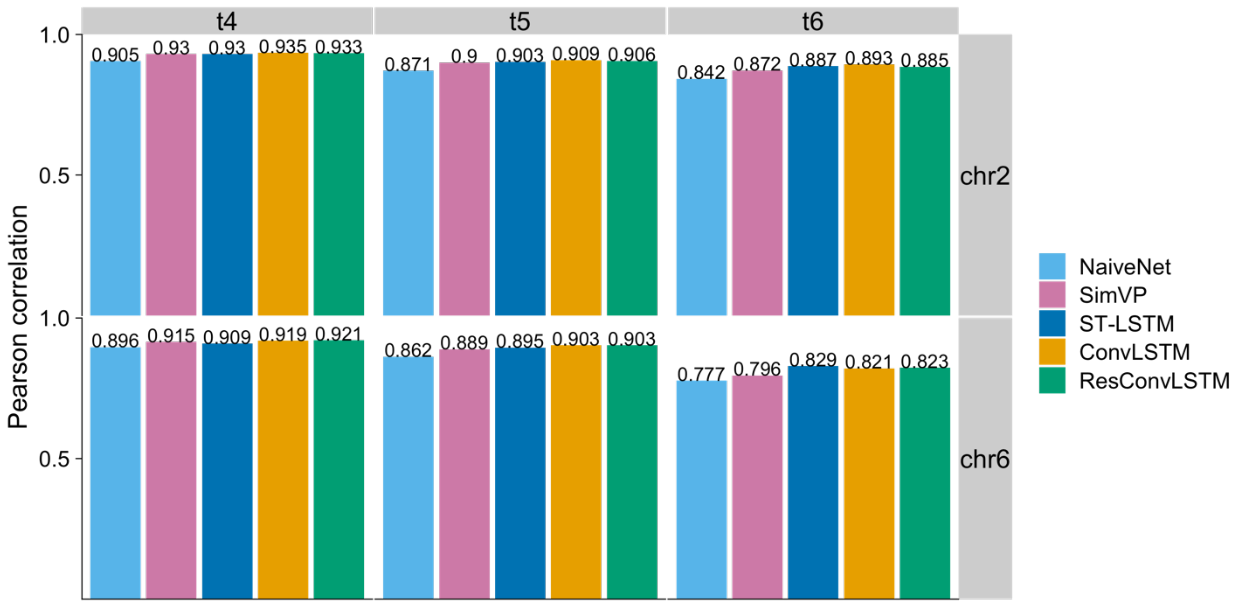


**Fig. S5.** Pearson correlations between insulation scores calculated on experimental (ground truth) and predicted Hi-C contact matrices on dataset 1.

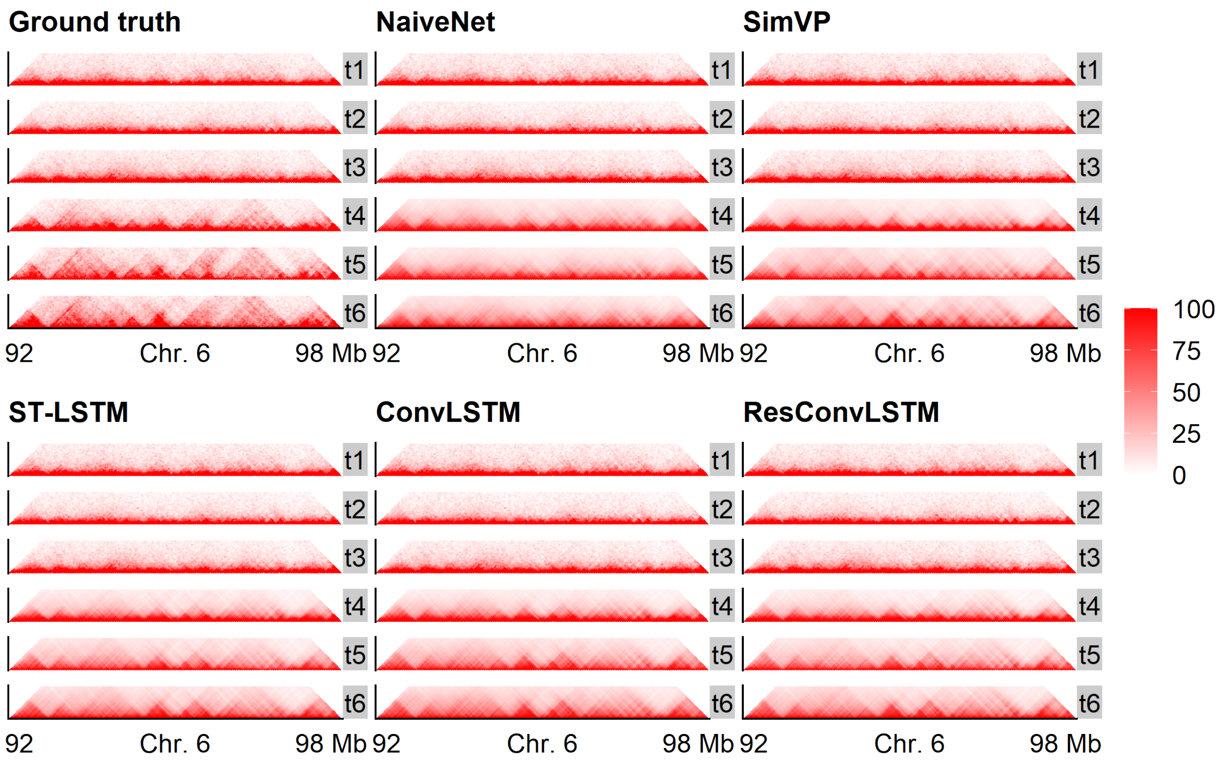


**Fig. S6.** Hi-C heat maps of ground truth and predictions from the five methods at the input ($t_1$, $t_2$, and $t_3$) and future ($t_4$, $t_5$, and $t_6$) time-steps.

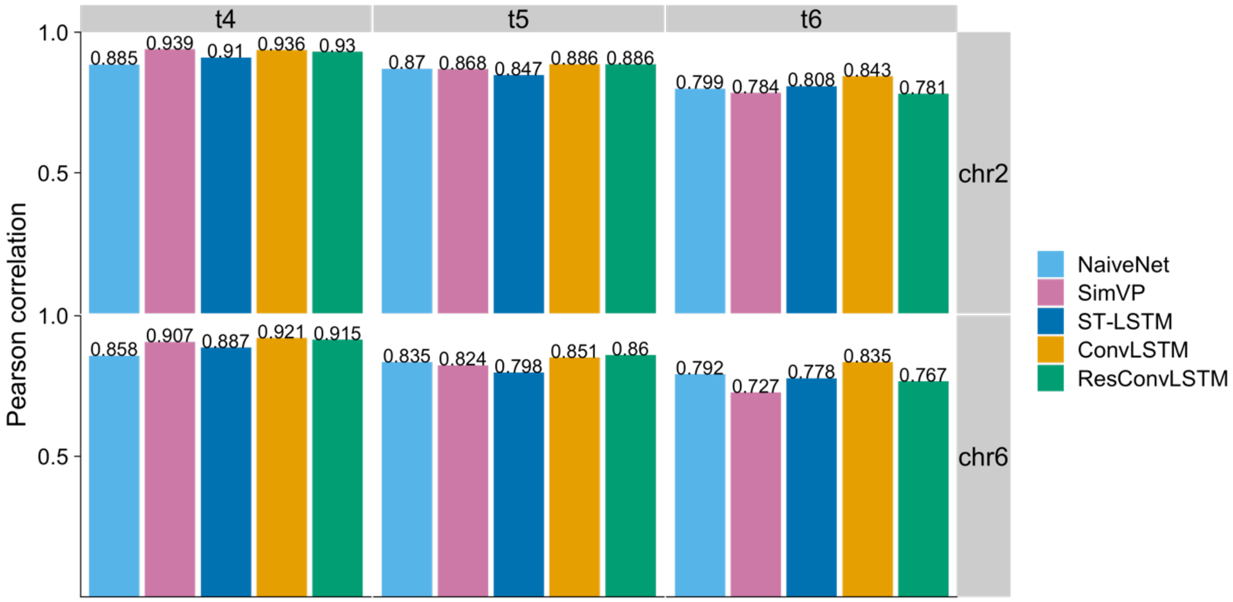


**Fig. S7.** Pearson correlations between insulation scores calculated on experimental (ground truth) and predicted Hi-C contact matrices on dataset 2.
